# A single microRNA-Hox gene module controls complex movement in morphologically-distinct developmental forms of *Drosophila*

**DOI:** 10.1101/511881

**Authors:** AR. Issa, J. Picao-Osorio, N. Rito, M.E. Chiappe, C.R. Alonso

## Abstract

Movement is the main output of the nervous system. It emerges during development to become a highly coordinated physiological process essential to the survival and adaptation of the organism to the environment. Similar movements can be observed in morphologically-distinct developmental stages of an organism, but it is currently unclear whether these movements have a common or diverse molecular basis. Here we explore this problem in *Drosophila* focusing on the roles played by the microRNA (miRNA) locus *miR-iab4/8* which was previously shown to be essential for the fruit fly larva to correct its orientation if turned upside down (self-righting) (Picao-Osorio et al., 2015). Our study shows that *miR-iab4* is required for normal self-righting across all three *Drosophila* larval stages. Unexpectedly, we also discover that this miRNA is essential for normal self-righting behaviour in the *Drosophila* adult, an organism with radically different morphological and neural constitution. Through the combination of gene-expression, optical imaging and quantitative behavioural approaches we provide evidence that *miR-iab4* exerts its effects on adult self-righting behaviour through repression of the *Hox* gene *Ultrabithorax (Ubx)* (Morgan, 1923; Sánchez-Herrero et al., 1985) in a specific set of motor neurons that innervate the adult *Drosophila* leg. Our results show that this *miRNA-Hox* module affects the function, rather than the morphology of motor neurons and indicate that post-developmental changes in *Hox* gene expression can modulate behavioural outputs in the adult. Altogether our work reveals that a common *miRNA-Hox* genetic module can control complex movement in morphologically-distinct organisms and describes a novel post-developmental role of the *Hox* genes in adult neural function.

## INTRODUCTION

Movement, the main output of the nervous system, first emerges during embryonic development. Although in its initial embryonic manifestation movement typically appears highly uncoordinated, as development proceeds, movement turns into an exceedingly coordinated physiological process (Landmesser and O’Donovan, 1984; Suzue, 1996; Saint-Amant and Drapeau, 1998; Crisp, 2008) that ultimately enables the fully formed animal to feed, escape from predators or find a suitable partner to mate. As such, adequate movement control is key to the animal’s adaptation to the environment and represents an essential intrinsic attribute fundamental to organismal survival and evolution. A point worth noting is that animals as distinct as insects and mammals all have a preferred orientation in respect for the surface so that their grip is maximised to facilitate motion.

To what extent does the genetic make-up of the organism influence the control of its movements? In principle, genetic mutations could affect the control of movement in two fundamentally distinct ways: they could impair the developmental formation of the networks underlying movement control, or, instead, interfere with the function of the cellular components involved in the physiological regulation of movement. A priori, these two levels of action of the genetic system need not be mutually exclusive.

A key experimental system to study the effects of genes on movement control is the fruit fly *Drosophila melanogaster*. Here, following the behavioural genetics approach pioneered by Seymour Benzer and his colleagues (Benzer, 1967; Hotta and Benzer, 1972), it became possible to isolate several genes with associated roles in movement control, including the Zinc-finger transcriptional co-repressor gene *scribbler* (Yang et al., 2000), the cGMP-dependent protein kinase gene *foraging* (de Belle et al., 1989; Osborne et al., 1997), the Ig superfamily gene *turtle* (Bodily et al., 2001), the phosphatidic acid transporter gene *slowmo* (Carhan et al., 2004) and other genes, such as *pokey* (Shaver et al., 2000) whose molecular functions have not yet been established. Of note is the case of the *Hox* genes, which encode a family of transcription factors key for the correct development of body structures along the main body axis (Lewis, 1978; McGinnis and Krumlauf, 1992; Alonso, 2002; Mallo and Alonso, 2013), and whose function has been shown to be required for the correct development of the neuromuscular networks underlying larval crawling (Dixit et al., 2008).

Nonetheless, much of the genetic dissection of movement control has so far focused on so-called protein-coding genes. Recent work in our laboratory showed that a single non-coding RNA, the microRNA *miR-iab4*, can affect the complex motor sequence that allows the young fruit fly larva to rectify its orientation if turned upside down (self-righting, SR) (Picao-Osorio et al., 2015). SR is an adaptive innate response that ensures an adequate position of the organism in respect for the surface, and it is evolutionarily conserved all the way from insects to mammals (Ashe, 1970; Penn, and Brockmann, 1995; Faisal. and Matheson, 2001; Jusufi, et al., 2011), including humans.

microRNAs (miRNAs) repress gene expression by blocking protein translation or promoting target mRNA degradation of their targets (Bartel, 2018). Our previous work showed that *miR-iab4* affects larval movement through regulation of one of its molecular targets, the *Hox* gene *Ultrabithorax* (*Ubx*), whose level of expression in a set of metameric motor neurons is critical for normal SR behaviour (Picao-Osorio et al., 2015). To explore the generality of the effects of miRNA regulation on larval SR movement we recently conducted a genetic screen that revealed that at least 40% of all miRNAs expressed in young *Drosophila* larva can affect SR demonstrating an unprecedented and widespread role of miRNA regulation in the control of postural adjustments and locomotor behaviour (Picao-Osorio et al., 2017).

Despite this progress, it is currently unclear whether similar (adaptive) movements performed by morphologically distinct organisms rely on common or different genetic operators. Here we investigate this problem looking at the effects of the *miR-iab4/Ubx* system on a series of distinct developmental stages of the fruit fly including the larvae and adults: organisms with substantially different somatic and neural anatomies, behavioural repertoire and life style.

Through the combination of gene expression, optical imaging and behavioural analyses we show that a single genetic module composed of the miRNA *miR-iab4* and the *Hox* gene *Ubx* contributes to the SR response in both *Drosophila* larvae and adults. Our study also reveals a novel neural role of the *Hox* genes in the fully formed adult, suggesting that these key developmental genes also perform biological functions once development has ceased.

## RESULTS AND DISCUSSION

Our previous work in the young, first instar *Drosophila* larvae showed that ablation of the *miR-iab4/8* locus (Bender, 2008) leads to significant defects in the SR response (Picao-Osorio et al., 2015). To investigate whether *miR-iab4/8-dependent* effects were confined to the L1 stage or had impact throughout larval development we conducted a series of SR tests in first, second and third instar larvae (L1, L2 and L3 larvae, respectively) (Figure 1A-B). SR was assayed by briefly putting individuals upside-down and monitoring the time they took to come back to a normal right side up position (see Materials and Methods). miRNA mutants show an increased time for the completion of the SR sequence (Figure 1C), indicating both that activities derived from the *miR-iab4/8* locus affect SR, and that this miRNA system is important for the normal timing of the SR response across all three larval stages.

**Figure 1.**
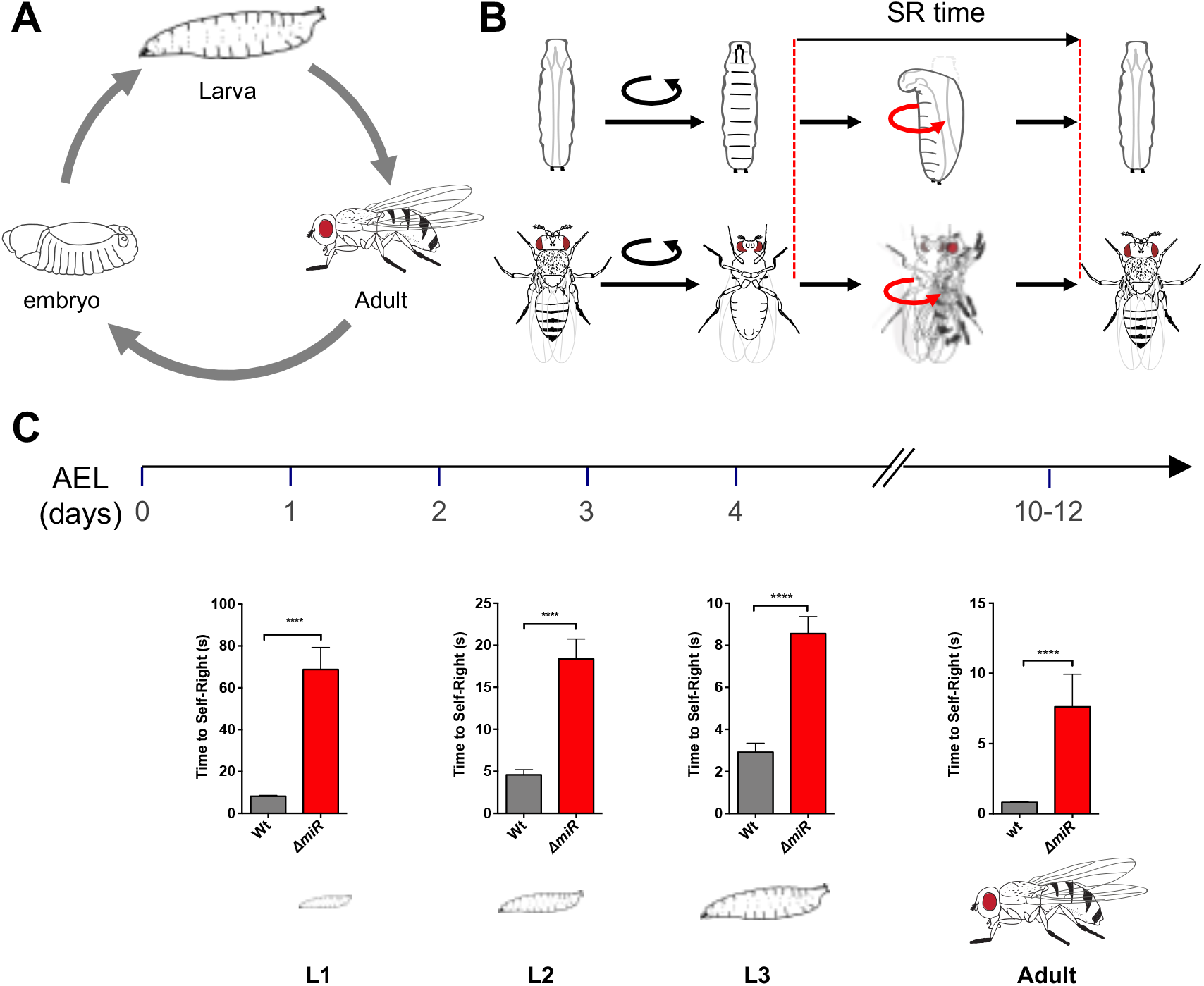
Removal of *miR-iab4/iab8* disrupts larval and adult self-righting behaviour (SR). **(A)** *Drosophila melanogaster* life cycle. **B.** Diagram of SR behavioural response in larvae (top) and adults (bottom). **(C)** Quantification of the time required for the successful completion of the SR behaviour along larval stages and in the adult (mean ± SEM; N = 63-70 flies for L1, 27-28 for L2, 25 for L3, and N= 49-54 adults) in wild-type controls (w^1118^, grey) and *miR-iab4/iab8* mutants *(ΔmiR*, red). Analysis of SR behaviour throughout development shows that *ΔmiR* mutants have defects across larval stages and in the adult. A nonparametric Mann-Whitney U test was performed to compare treatments; p < 0.001 (***).

Like in all holometabolous insects, the *Drosophila* life cycle involves the transformation of the larva into the adult through the process of metamorphosis (Truman and Riddiford, 1999). Given the substantial anatomical and functional remodelling that metamorphosis imposes on the *Drosophila* soma and nervous system, genetically-induced behavioural defects observed in the larvae may simply disappear in the adult stage. Remarkably, a modification of the SR test performed in the *Drosophila* adult (see Materials and Methods), reveals that integrity of the *miR-iab4/8* locus is important for normal SR response also in the adult fly (Figure 1C and Figure S1). That is, a common miRNA system controls the same adaptive behaviour in two stages of the life of a fly, with radically different morphological and neural constitution.

Importantly, analysis of free-walking behaviour in adult flies (see Methods) show that the mutation of the *miR-iab4/8* locus does not impair locomotion in adult flies (Figure S2). This indicates that the absence of the *miR-iab4/8* system does not lead to a generalised motor deficient phenotype. However, miRNA mutant flies tend to walk less than controls, and previous work (Bender, 2008) showed that the *miR-iab4/8* mutants displayed posture control defects during mating, suggesting that *miR-iab4/8* may function in other posture control systems in addition to SR. Alternatively, these different posture control systems may share some of the same neural substrate upon which *miR-iab4/8* exerts its biological role.

The *miR-iab4/8* locus encodes two distinct miRNA molecules: *miR-iab4* (Ronshaugen et al., 2005) and *miR-iab8* (Bender, 2008; Tyler et al., 2008; Stark et al., 2008), each produced from pri-miRNA transcription of opposite DNA strands. To tease apart the individual contributions of *miR-iab4* and *miR-iab8* towards adult SR we performed a series of genetic tests in adults using a collection of chromosomal variants that specifically disabled *miR-iab4* or *miR-iab8* placing them in combination with the *miR-iab4/8* mutation *(DmiR)* and determined that *miR-iab4* (and not *miR-iab8*) is responsible for the effects on the adult SR response (Figure S3).

Behavioural observation of the SR routine in the adult shows that legs perform a key role during the SR response (Figure 2A), allowing the animal to swiftly grab the substrate and use this point of contact to flip its body into the right side up position. The halteres, important mechanosensory organs that control body manoeuvres in flight (Nalbach, 1993), may also contribute to the control of body manoeuvres underlying the SR response. However, we found that flies with ablated halteres showed no effect in the time to complete the SR response as compare to controls (Figure S1C). Next, we asked which pair of legs derived from the pro-(T1), meso- (T2), or meta- (T3) thoracic segments contributed to the control of SR. We performed a series of ablation experiments in which we surgically removed T1-, T2-or T3-legs from wild type individuals and assessed their performance in SR tests (Figure 2B). These experiments showed that the activities of T1- and T3-legs contribute to normal SR, whereas removal of T2-legs had no detectable effects.

**Figure 2.**
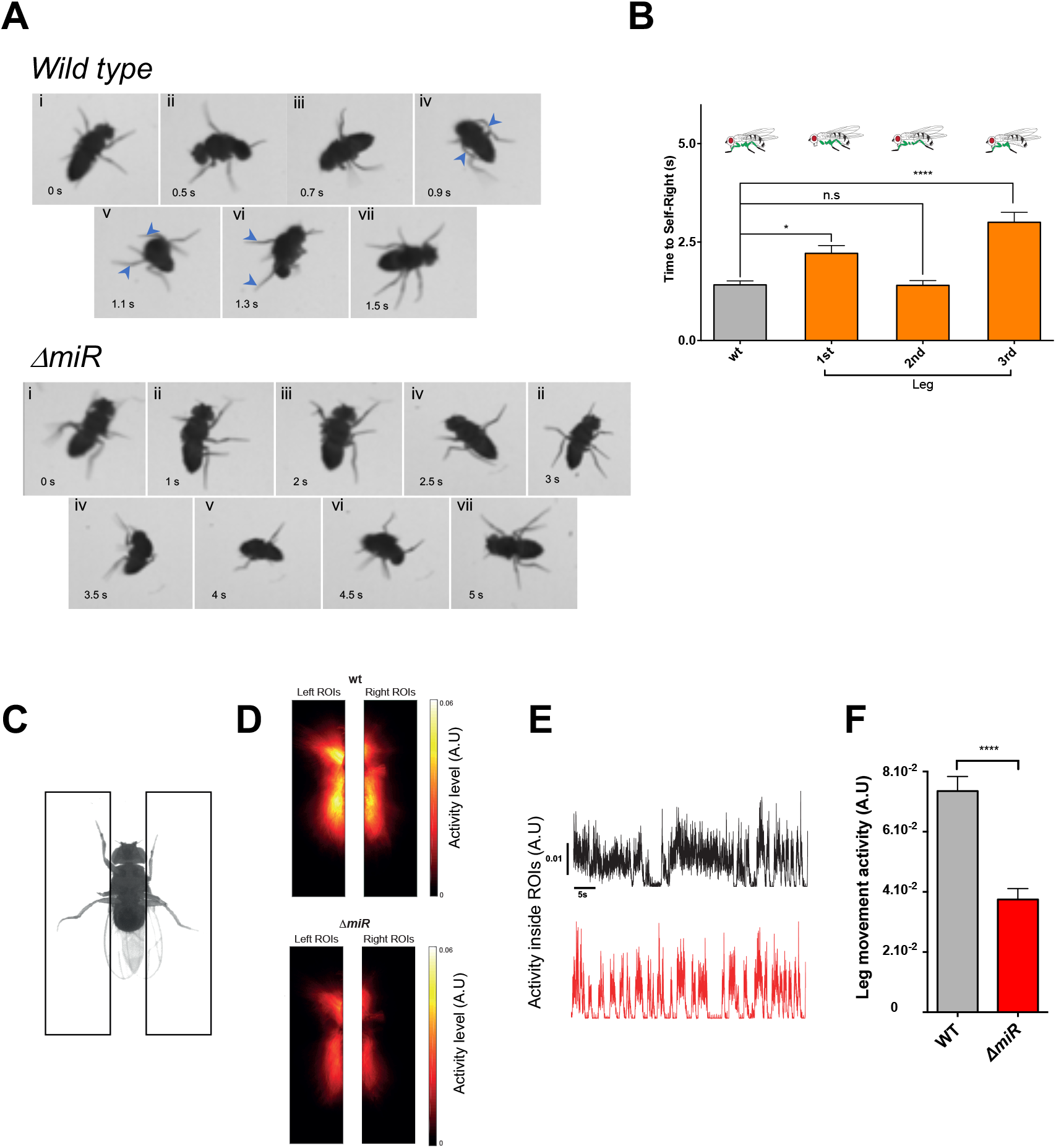
Analysis of leg activity levels reveal the effect of *ΔmiR* on the control of the SR response. **(A)** Time-lapse sequences of the SR response in WT and *ΔmiR* flies (i. Upside down position, ii. Movement of legs, iii. Fixing of 3rd leg on the substrate, iv. Initiation of the tilting, v. Standing position, vi. Whole body tilting, vii. Back to the normal position). **(B)** Quantification of the time to complete the SR response in WT flies without T1, T2 and T3 legs (pair) (mean ± SEM; N = 40 flies). **(C-F)** Quantification of leg activity levels in WT control (w^1118^) and *miR-iab4/iab8* mutant *(ΔmiR)* flies. **(C)** Schematic of the regions of interest (black rectangles) used to quantify leg activity levels (see Materials and Methods). **(D)** Average heat maps of leg activity for WT control (top), and *ΔmiR* mutant (bottom) flies. Color code indicates amplitude of activity, with warm colors representing high levels. **(E)** Average leg activity level (mean ± SEM; N = 14 flies for control and *ΔmiR* flies). A nonparametric Mann-Whitney U test was performed to compare groups; p < 0.0001 (****).

We then sought to establish whether leg movement showed any anomalies in *miR-iab4/8* mutant flies when compared with wild type specimens. Quantification of leg movement in immobilised upside-down flies showed that in *DmiR* mutant flies legs displayed a reduction in activity levels compared to those observed in WT flies (Figure 2D-E,). This observation suggests that the impact of *miR-iab4* on the SR response is mediated –at least in part– through effects in the levels of activity of the legs (Figure 2F-G).

To explore the molecular basis underlying *miR-iab4* effects on adult SR we considered the hypothesis that *miR-iab4* exerts its effects on adult SR via the same molecular target established in the larva, the *Hox* gene *Ubx* (Picao-Osorio et al., 2015) (Figure 3A). In addition, *Ubx* plays a key developmental role in allocating morphological specificity to one of the thoracic ganglia (T3) (Lewis, 1978; Mallo and Alonso, 2013), including effects on detailed patterning of the T3-leg (Rozowski and Akam, 2002). To test the model that *miR-iab4* represses *Ubx* to allow for normal adult SR response, we increased the expression levels of *Ubx* within its natural transcriptional domain in normal flies seeking to emulate the de-repression effects caused by miRNA removal. The results of this experiment (Figure 3B) show that an increase of *Ubx* levels phenocopies the effects of the *miR-iab4/8* mutation on adult SR response, suggesting that the expression levels of *Ubx* are important for normal behavior.

**Figure 3.**
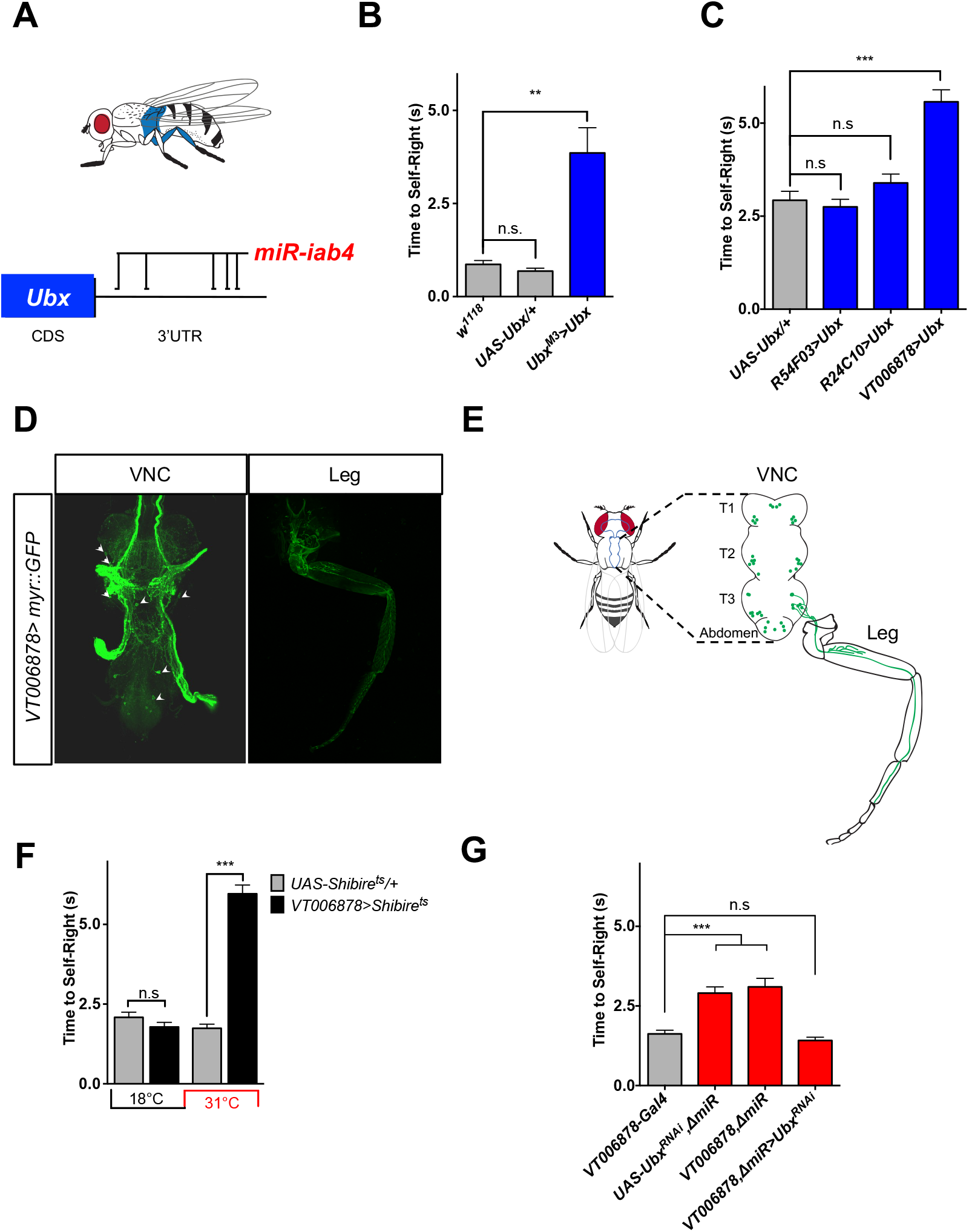
miRNA-dependent *Ubx* regulation in ventral VT006878/lin15 motor neurons underlies adult SR behaviour. Roles of specific motor neuron subpopulations in SR behaviour. **A**. The *Hox* gene *Ubx* is expressed in the third thoracic region (blue); previous work (see main text) showed that *miR-iab4* regulates *Ubx* expression via specific target sites in *Ubx* 3’UTR sequences. **B**. Quantification of SR behaviour in adult flies overexpressing *Ubx* within its natural expression domain (Ubx^M3^>Ubx) shows that upregulation of *Ubx* is sufficient to cause an adult SR defect (mean ± SEM; n = 19-25). **C**. *Ubx* overexpression in the VT006878/lin15 motor neurons innervating T3 legs phenocopies SR abnormal response (mean ± SEM; N = 27-28 flies). **D**. Diagram describing the pattern of VT006878-Gal4 expression in the adult VNC and T3 leg. **E**. Confocal images of *VT006878>GFP* in the VNC (left) and leg (right); arrows show cell bodies. **F**. Blocking neural activity in VT006878 neurons leads to defects in adult SR response (mean ± SEM; N = 57-64 flies). **G**. In *ΔmiR* flies, RNAi-mediated decrease of *Ubx* expression within the VT006878 domain rescues the SR phenotype (mean ± SEM; N = 41 flies). A nonparametric Mann-Whitney U (Figure A, B, F) and one-way ANOVA with the post hoc Tukey-Kramer (Figure G) tests were performed to compare treatments; p > 0.05 (nonsignificant; n.s.), p < 0.05 (*) and p < 0.001 (***).

Taking into consideration: (i) that mutation of the *miR-iab4/8* locus disrupts the SR response; (ii) that legs play a key role in the SR sequence, and (iii) that modulation in the levels of the *miR-iab4* target *Ubx* within its transcriptional domain had significant impact on SR, we decided to explore the cellular basis underlying SR control by testing the model that modulation of *Ubx* in leg motor neurons – the direct effectors of leg activity – may play a role in the adult SR response. For this, we artificially upregulated *Ubx* in different neuronal assemblies known to innervate the *Drosophila* leg (Bierley, 2012; Baek and Mann, 2009; Lacin and Truman, 2016) using the available lineage specific leg motor neuron GAL4 drivers *VT006878-GAL4* (NB2-3/lin15) and *R24C10-Gal4* (NB5-7/lin20). These experiments showed that upregulation of *Ubx* within the domain demarked by the *VT006878-Gal4* (Lacin and Truman, 2016) was sufficient to cause an increase in the time that individual flies take to complete the SR response (Figure 3C and Figure S5), whereas the other motor neuronal drivers had no effect. In addition, inactivation of *VT006878* neurons (Figure 3D-E) through expression of a temperature-sensitive allele of *shibire* (Kitamoto, 2001) has pervasive effects on the timing of the SR response (Figure 3F), highlighting the contribution of these motor neurons to the normal SR response.

We noted that the *VT006878* line seems to be transcriptionally active in wing and haltere sensory axons (Figure 3D), making it plausible that wings and/or halteres may play a role in adult SR. However, different manipulations indicated this might not be the case. First, all adult flies had surgically removed wings in our SR behavioural paradigm, making it unlikely that these appendages are key contributors to this behaviour. Second, ablation of halteres resulted in no apparent effect in the time wild type flies took to complete a SR response (Figure S1C). Altogether, these results suggest that the effect observed by the overexpression of *Ubx* in the *VT006878-Gal4* line cannot be accounted by an effect derived from the haltere/wing sensory neurons.

Upregulation of *Ubx* using *VT006878-Gal4* is expected to increase *Ubx* levels in all three thoracic segments (T1-T3) and scattered neurons in the brain (Lacin and Truman, 2016), making it unclear whether ectopic *Ubx* expression in the brain might be the cause of the SR defects observed in treated adults. To constrain the expression pattern of *VT006878>Ubx* to the brain only, we used the ventral nerve cord (VNC) specific tool *teashirt-Gal80^ts^*. Upregulation of *Ubx* within circuits in the brain has no effect on the timing of SR in adults (Figure S4) indicating that: (i) an increase of *Ubx* within the *VT006878* domain in the brain is insufficient to cause an adult SR phenotype; and (ii) *Ubx* upregulation within the thoracic *VT006878* domain is indeed responsible for the triggering of SR defects in the adult.

Ubx protein is detected in subsets of neurons within the T1-T3 ganglia, with a larger population observed within the T3 segment of the VNC (Figure 4A-B). The RNA *in situ* hybridisations show that *miR-iab4* is highly expressed in the T3 ganglion of the VNC (Figure 4A and C, E-F) and that both Ubx and miR-iab4 are expressed within the *VT006878* domain (Figure 4D and G). In miRNA mutants, *Ubx* expression is significantly increased in the T3 segment of the VNC, but not in T2 (Figure 4H-I and Figure S6), in agreement with the idea that increase of *Ubx* expression (de-repression) within the *VT006878* domain in T3 leads to SR defects in the adult. A prediction that emerges from this idea is that artificial reduction of *Ubx* in *DmiR* mutants, specifically confined to the *VT006878* domain should ameliorate (or even rescue) the SR phenotype observed in adult mutants. In line with this prediction, RNAi-mediated reduction of *Ubx* driven by *VT006878-Gal4* rescues the SR phenotype in adult flies (Figure 3G). This experiment also indicates that levels of *Ubx* in the T3 segment are key for normal timing of adult SR.

**Figure 4.**
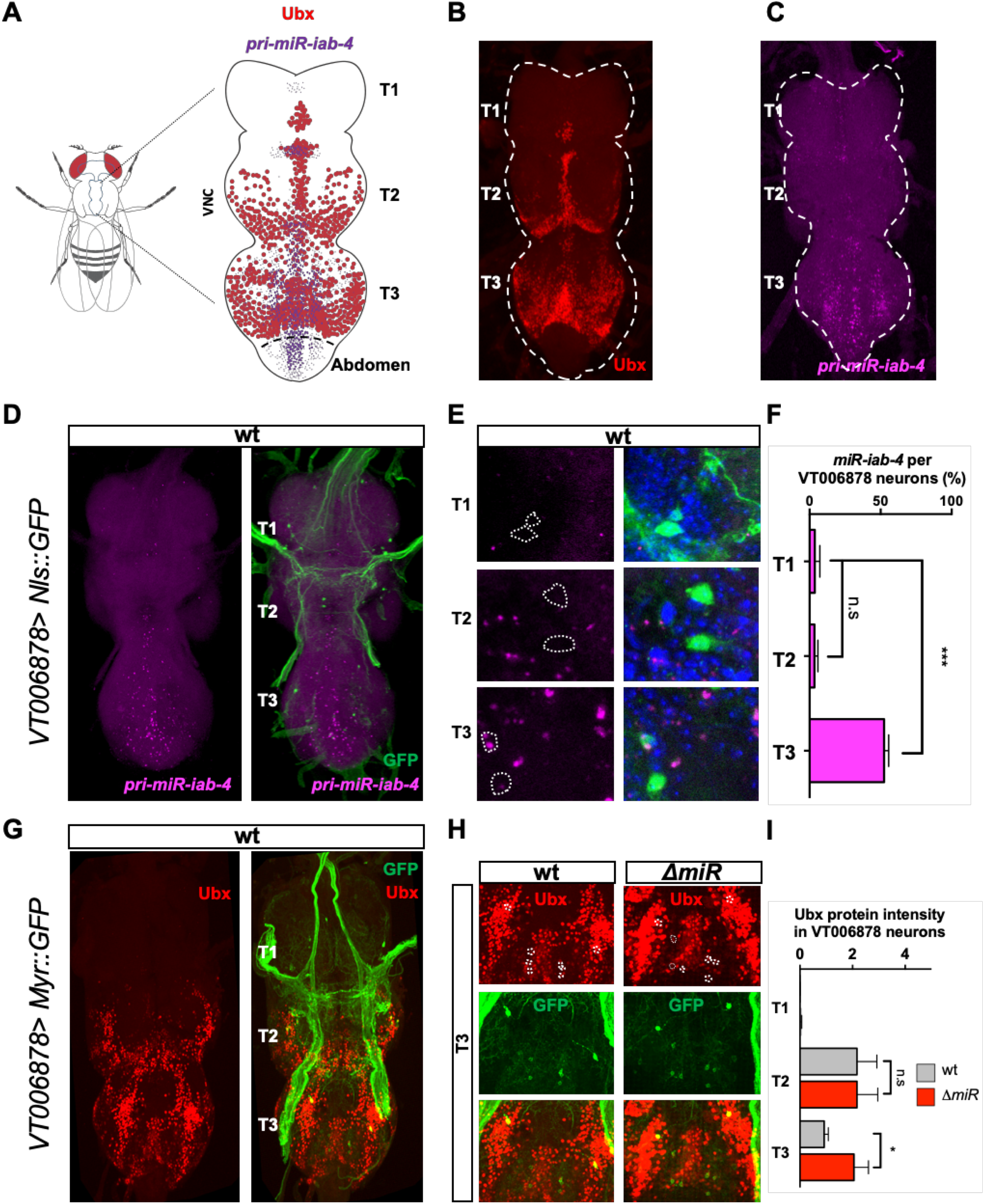
*Expression analysis of* Ubx protein and *miR-iab-4* in the adult VNC. **A**. Schematic diagram of the adult VNC showing areas of *miR*-*iab-4* (magenta) and Ubx protein (red) expression. **B**. Expression of Ubx protein within the VNC shows highest signal in the T3 ganglion, but signal is also detectable in T2 and a much lower level in T1. **C**. *miR-iab-4* expression in the VNC shows highest level of expression in the T3 ganglion. **D**. *miR-iab-4* expression profile in VT006878 positive neurons. **E-F**. Quantification of *miR-iab4* signal (magenta) within the VT006878 domain (green) along different ganglia of the VNC shows a significant increase of miRNA signal in the T3 ganglion (Blue: DAPI) (mean ± SEM; N = 7 VNC). **G**. Expression of Ubx protein (red) is detected within the VT006878 domain (labelled by GFP, green). **H**. Expression pattern of Ubx protein within the VT006878 domain in the T3 region of the VNC in wild type and miRNA mutants. A significant increase in Ubx protein expression is observed in the T3 ganglion of mutant adult flies; in contrast, comparison of normal and miRNA mutant flies shows no differences in Ubx expression in the T2 ganglion (mean ± SEM; n = 7-13 VNC). A nonparametric Mann-Whitney U and one-way ANOVA with the post hoc Tukey-Kramer tests were performed to compare treatments; P > 0.05 (nonsignificant; n.s.), p < 0.05 (*) and p < 0.001 (***).

Detailed anatomical examination of T3 *VT006878* motor neurons (also known as *ventral lineage 15 motor neurons* (Lacin and Truman, 2016)) in wild type and *DmiR* specimens showed no detectable differences in axonal projections or morphologies (Figure 5A-C) suggesting that – as observed in the larva (Picao-Osorio et al., 2015) – the microRNA under study might have effects on neuronal function rather than on neuronal morphology. Indeed, quantification of varicosities (a known indicator of neuronal activity (Cox et al., 2000; Petreanu et al., 2012) at the junction of VT006878 neurons with the muscle system of the third leg reveals a statistically-significant reduction in varicosities in the *DmiR* samples (Figure 5D-E) in line with the model that absence of the miRNA leads to diminished levels of neural activity. Remarkably, in *DmiR* mutants, RNAi-mediated reduction of *Ubx* expression in the VT006878 neurons rescues the normal number of varicosities strongly indicating a role of Ubx in the formation of active contact points between the neuronal and muscle systems. Furthermore, multiphoton microscopy analysis of genetically-encoded calcium reporters (GCaMP6m) specifically expressed in the VT006878 motor neurons shows an overall reduction of spontaneous neural activity in *DmiR* samples in T3 when compared to wild type (Figure 5F-G). Altogether, our data suggests that *miR-iab4* represses *Ubx* within the VT006878 motorneuron domain in T3 ensuring the normal neural functions that underlies the adult SR response.

**Figure 5.**
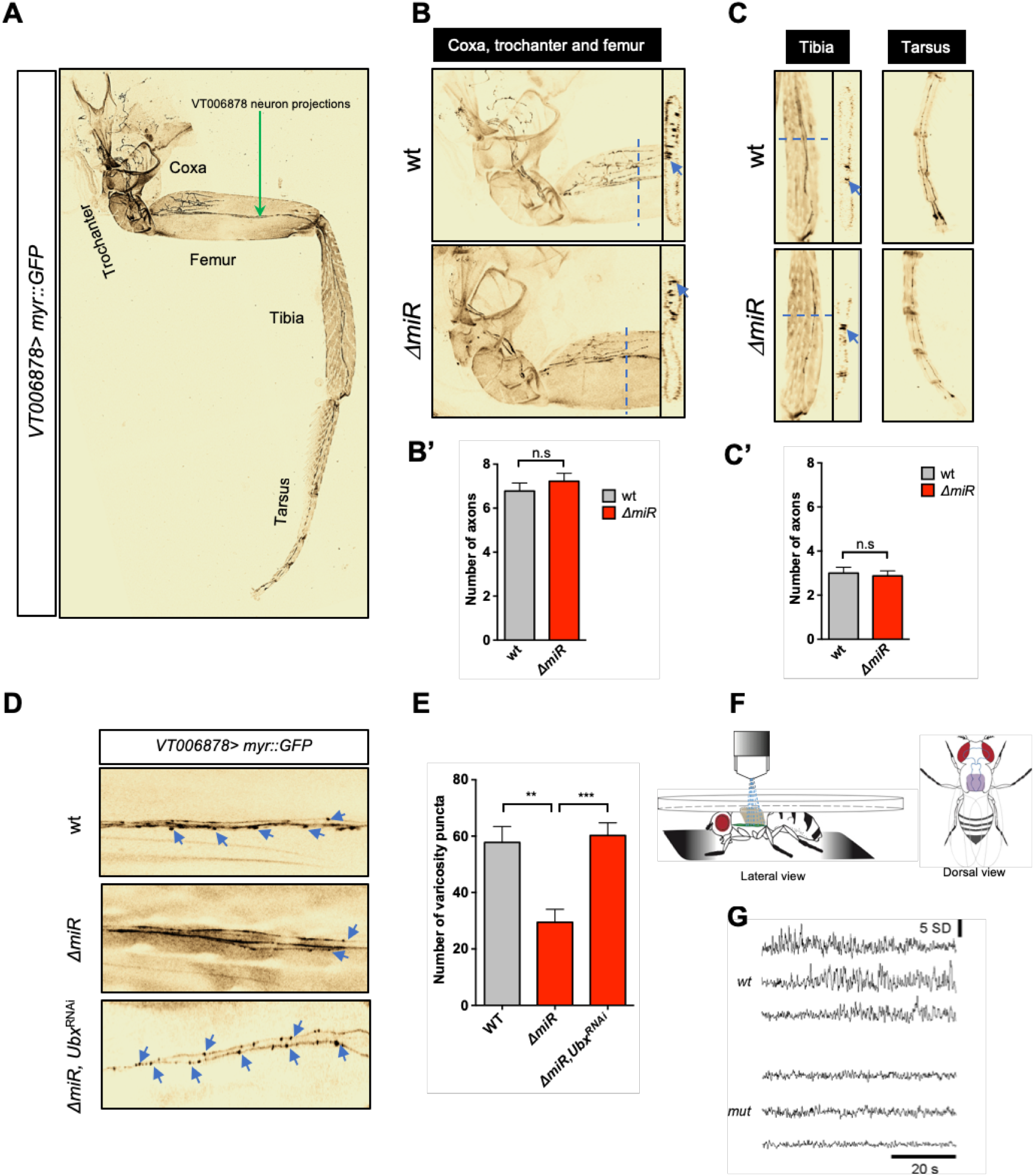
Effects of miRNA mutation on the morphology and function of VT006878/lin15 motor neurons. **A**. Image of hind (T3) leg showing VT006878 positive neuronal projections labelled by GFP *(VT006878>Myr::GFP).* VT006878 neurons innervate the coxa, trochanter, femur, tibia and tarsus segments. **B, B’**. Quantification of projections of VT006878 neurons into the coxa, trochanter and femur of wt and *DmiR* specimens shows no differences across genotypes (mean ± SEM; n = 9). **C, C’**. Analysis of VT006878 projections in the tibia and tarsus shows no effects of the miRNA system on VT006878/lin15 morphology (Arrows highlight some of the motor neuron projections analysed) (mean ± SEM; N = 8 flies). **D, E**. Varicosity puncta of VT006878 projections in wt and *DmiR* femur. The area where these images were taken are indicated by the yellow dashed rectangle on panel A. Note the significant reduction in puncta observed in miRNA mutants and the dramatic effects caused by Ubx RNAi treatment within the VT006878 domain in miRNA mutants which rescues the normal number of puncta as observed in wild type samples (mean ± SEM; N = 10-12 flies). **F**. Schematic representing the preparation used for neural activity recordings. **G**. Representative neural activity traces with time indicated by standard normalised fluorescence (SD) of wt (top) and *ΔmiR* (bottom). A nonparametric Mann-Whitney U (Figure B’ and C’) and one-way ANOVA with the post hoc Tukey-Kramer (Figure E) tests were performed to compare treatments; p > 0.05 (nonsignificant; n.s.), p < 0.01 (**) and p < 0.001 (***).

Lastly, we sought to determine whether the effects of the miRNA on adult SR behaviour emerge from a progressive developmental function of the miRNA, or rather, are the consequence of the activity of the miRNA on the physiology of the *VT006878* motor neurons in the adult. For this we performed a conditional expression experiment in which we maintained normal expression of *Ubx* in the VT006878 domain during the full developmental process that spans from embryo to adulthood, increasing *Ubx* expression only after adult eclosion (Figure 6A). Our data shows that an increase in *Ubx* expression, exclusively delivered in the adult, is *sufficient* to induce SR defects (Figure 6B and Figure S7) revealing a novel post-developmental role of *Hox* genes in the control of neural function in the fully formed organism.

**Figure 6.**
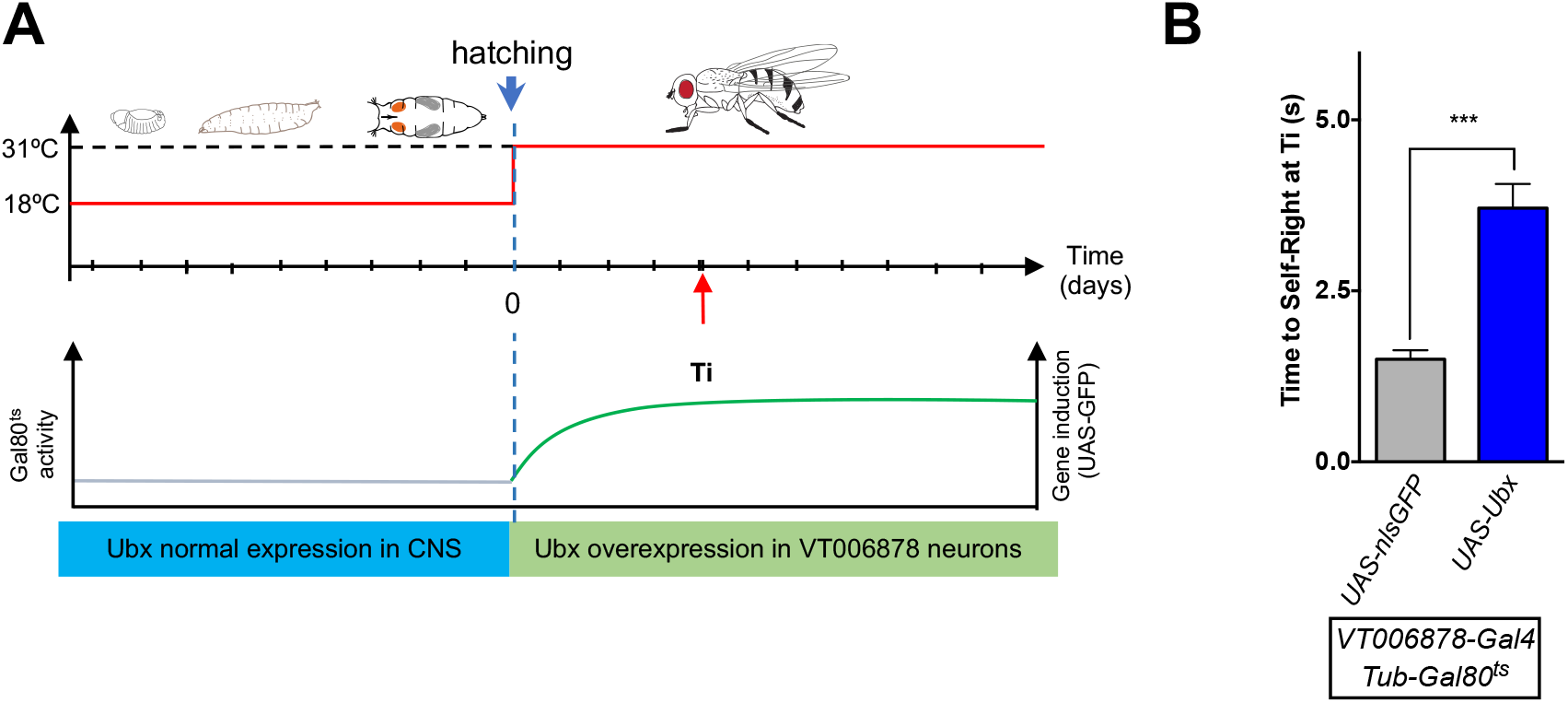
Conditional increase of *Ubx* expression after development. **A**. Conditional expression experiment in which Ubx protein is upregulated only once development has been completed. Graphic representation of *Gal4* and *Gal80* activities over developmental time. N.B. At 18°C, *Gal80^ts^* represses Gal4 activity; at 31°C, the *Gal80^ts^* role is inactivated allowing for *VT006878-Gal4* mediated induction of Ubx (green) in lin15 neurons. Maximal induction is achieved approximately 4-days after eclosion (Ti). **B**. SR behaviour quantification of flies at Ti reveals that post-developmental induction of Ubx in VT006878 neurons is sufficient to cause SR defects (mean ± SEM; N = 19-25 flies). A nonparametric Mann-Whitney U test was performed to compare treatments; p < 0.001 (***).

Our work reveals that complex adaptive movements performed by organisms with distinct morphologies and neural anatomies can rely on a simple genetic module involving a miRNA and a *Hox* gene. We also describe what is –to our best knowledge– the first case of post-developmental roles of the *Hox* genes with impact on adult behaviour. Based on the pervasive evolutionary conservation on the *Hox* gene system and the key roles played by these genes in the nervous systems of animals as different as insects and mammals, we propose that similarly simple genetic modules including miRNAs and *Hox* targets may conform part of the molecular circuitry underlying movement control in other species, including humans.

## ACKNOWLEDGEMENTS

We thank members of the Alonso lab for helpful discussions and comments. We wish to thank Filip Janiak and Tom Baden (University of Sussex) for their assistance with multiphoton microscopy experiments and Clare Hancock for preliminary observations in this project. This research was funded by a Wellcome Trust Investigator Award made to C.R.A. (098410/Z/12/Z).

## MATERIALS AND METHODS

### Drosophila Culture and Strains

Fly stocks and crosses were raised at 25°C on standard corn meal/yeast/agar medium, under a 12 h/12 hr light/dark cycle. The following fly strains were used: *VT006878-Gal4* (Vienna Drosophila Resource Center, ID200694); *R54F03-Gal4* (BDSC, #39078); *R24C10-Gal4* (BDSC, #49075) (Lacin and Truman, 2016); *ΔmiR-iab4/iab8* (Bender, 2008); iab-3^277^; iab-5^105^ and *iab-7MX2* (Karch et al., 1985); *UbxM3-Gal4* (De Navas et al., 2006); *UAS-UbxIa* (Reed et al., 2010); *UAS-UbxRNAi* (BDSC, #31913); *UAS-Myr::GFP* (Pfeiffer et al., 2010); *UAS-Nls*::GFP(BDSC, #4775); UAS-GCaMP6m (Chen et al., 2013); UAS-shi^ts^ (Kitamoto, 2001).

### Self-Righting tests

Larval SR behaviour was assayed as previously described (Picao-Osorio et al., 2015; Picao-Osorio et al., 2017). For adult SR behaviour tests, flies were grown in non-crowed conditions at 25°C. The day before the SR test, the wings of cold-anesthetised 2-to-4-day old flies were surgically removed (clipped). Flies recovered for one day at 25°C. Flies were assayed for SR behaviour by being introduced individually into an arena and rolled over with a brush to an “upside-down” position (“legs up”) and the time taken by the fly to return to its normal position (“right-side up”) was recorded. A maximum of ten minutes was given to the fly to SR. All experiments were done with flies 4-6 days after eclosion and tested at 25°C. For silencing of VT006878 neurons, 3-to 4-day-old flies expressing shi^ts1^, were incubated for 10 min at 31°C, or at 18°C for controls, just before the SR test. SR behaviour was assessed within seconds (50±10) after incubation. Similar results to those observed using this procedure were obtained when measuring the time to SR of adult flies with intact wings after recovery from CO_2_-induced or cold-induced anaesthesia (Figure S1). The absence of halteres showed no effects on SR (Figure S1).

### Walking behaviour

Locomotion was assessed by placing single flies (males or females) on a circular arena with sloped edges (Simon and Dickinson, 2010), and their spontaneous walking behaviour was recorded from the top at 60 Hz with a Flea FL3-U3-32S2M Point Grey camera with a M1214-MP2 lens (Computar) for 15 minutes. To automatically track the position of flies in the arena, we used Ctrax (Branson et al., 2009). Walking bouts were defined as segments in time when the body moved through space with speed of at least 5 mm/s for (at least) 500ms. As a measurement of locomotion performance, we calculated the straightness of a walking bout. Straightness was defined as the mean angular deviation from a line defined between the start and end points of the segment. A straightness of 0 indicated walking along a perfect straight line. Straightness greater than 0 indicated curvilinear trajectories, and the greater the value, the more prominent the deviations were from a straight course.

### Quantification of leg movements

Flies 3-5 days post eclosion, were cold-anesthetised and its thorax tethered to a glass microscope slides with UV-activated glue (BONDIC^®^). We next removed the tibia of mid leg to avoid interference of this leg in the quantification of first and third leg movements (this leg had no apparent effect on the timing of the SR response, Figure 2B). The leg movement videos were captured at 200 Hz with a monochrome digital camera (Bonito CL-400B, Allied Vision, with a M1214-MP2 lens and EX2C extender from Computar). We used a custom-made MATLAB script to quantify leg activity levels. Regions of interests (ROI) for analysis were automatically drawn based on the centre of mass (CM) of the thorax of the fly. Video images were converted into binary values using a threshold, and a time averaged image was calculated. Because the thorax of the fly was glued to a cover slip it was the only part of the fly that remained stationary throughout the video. To isolate pixels corresponding to the fly thorax, we identified those that did not change in intensity for more than 90% of the video. Pixels that did were converted to a background-related pixel. Next the thorax of the fly and its CM, were extracted using the connected components method. From the CM, we automatically defined two regions of interest (ROIs), one on each side of the fly that were separated by the width of the fly thorax, and with 459×100 pixel size. For each pixel inside of these ROIs, we extracted the pixel intensity (in A.U) and calculated the change in pixel intensity as a function of time. Leg activity per pixel was classified as 1 if the instantaneous change in pixel intensity was at least 10 pixels per time step, which corresponded to approximated 20%of the total change in pixel intensity. From this, we averaged the change in pixel intensity over the course of the experiment and generated a mean heat map (over all flies) for WT and *miRiab4/8* flies (Figure 2D). To quantify the average leg activity for each fly (Figure 2E), we calculated the mean response of both the Left and Right ROIs, normalized by the area of each ROI.

### Adult leg preparation and mounting

Tissue dissection and mounting were performed as described (Enriquez et al., 2016). Fly legs were dissected with forceps in 0.3% triton in 1x phosphate buffered saline (PBS). Adult legs attached to thoracic segments were fixed in 4% formaldehyde in PBS overnight at 4°C followed by five washes in PBTx for 20 minutes at room temperature. legs were mounted onto glass slides using 70% glycerol medium.

### Immunohistochemistry and RNA *in situ* hybridisation

Adult brains were dissected in 1X PBS. Tissues were then fixed for 1h in 4% formaldehyde in 1X PBS at room temperature. After fixation, brains were washed 3 times (30 min per washing) in PBS with 0.3% Triton-X-100 (PBTx) and incubated at 4°C overnight in primary antibodies. The following primary antibodies were used: mouse monoclonal anti-Ubx (FP3.38 (White and Wilcox, 1985) 1:500 from the Developmental Studies Hybridoma Bank) and chicken anti-GFP (Abacam Probes, 1:3000). The secondary antibodies were anti-mouse Alexa Fluor 555 (Invitrogen Molecular Probes, 1:1000) and anti-chicken Alexa Fluor 488 (Invitrogen Molecular Probes, 1:1000). RNA *in situ* hybridisation in adult ventral nerve cords for the precursor RNA transcripts of *miR-iab-4* was performed by designing 48 unique 20nt-probes labelled with Quasar 570 in the Stellaris platform from Biosearch Technologies, and using an adapted version of the protocol by Raj A. and Tayagi S., 2010. Images were acquired with a Leica SP8 confocal microscope, processed and analysed using FIJI Image J. The VT006878 nerve-ending varicosities were quantified by measuring the puncta they covered in VT006878-labelled by myrGFP in leg or VNC.

### Two-photon calcium imaging

To prepare flies for *in vivo* imaging in the ventral nerve cord (VNC) (Figure 5F) we adapted existing methods (Chen et al., 2018; Seeholzer et al., 2018). In brief, a single fly (3–5 days old) was cold-anaesthetised after eclosion and tethered using UV-curable glue to a piece of aluminium foil that covered a hole in the bottom of a modified polystyrene weighing dish. The fly’s body was positioned such that the dorsal side of the thorax covered the small hole made in the centre of the aluminium foil. The dish was then held by blue-tack on a glass microscope slide with ventral side and legs facing the slide. Next, the dish was filled with saline solution and a small hole in the thorax was opened by removing the cuticle covering the T3 ganglion using sharp forceps to avoid damaging nerves. The preparation was positioned under the two-photon microscope (see details below) and spontaneous GCaMP6m activity within VT006878 neurons was recorded. Composition of saline solution was as used previously (Seeholzer et al., 2018): 108 mM NaCl, 5 mM KCl, 2 mM CaCL_2_, 8.2 mM MgCL_2_, 4 mM NaHCO_3_, 1 mM NaH_2_PO_4_, 5 mM trehalose, 10 mM sucrose, 5 mM HEPES pH 7.5. All imaging experiments were performed on a MOM-type two-photon microscope (designed by W. Denk, MPI, Martinsried; purchased from Sutter Instruments/Science Products) equipped with a mode-locked Ti:Sapphire, Chameleon Vision-S laser set at 927nm. Emitted fluorescence was detected with F48×573, AHF/Chroma, and a water immersion objective 20x/1,0 DIC M27 Zeiss was used for image acquisition. For image collection we used custom-made software running under IGOR pro 6.3 for Windows (Wavemetrics) (Zimmermann et al., 2018), at 64 × 32 pixel resolution with 15.625 frames per second image sequences for activity scans or 512 × 512 pixel images for high-resolution morphology scans.

### Statistical analysis

Statistical analyses were performed with GraphPad Software Prism using Mann-Whitney U test or one-way ANOVA with the post hoc Tukey-Kramer test. Error bars in figures represent SEM. Significant values in all figures: *p < 0.05, **p < 0.01, ***p < 0.001.

---

**Figure S1.**
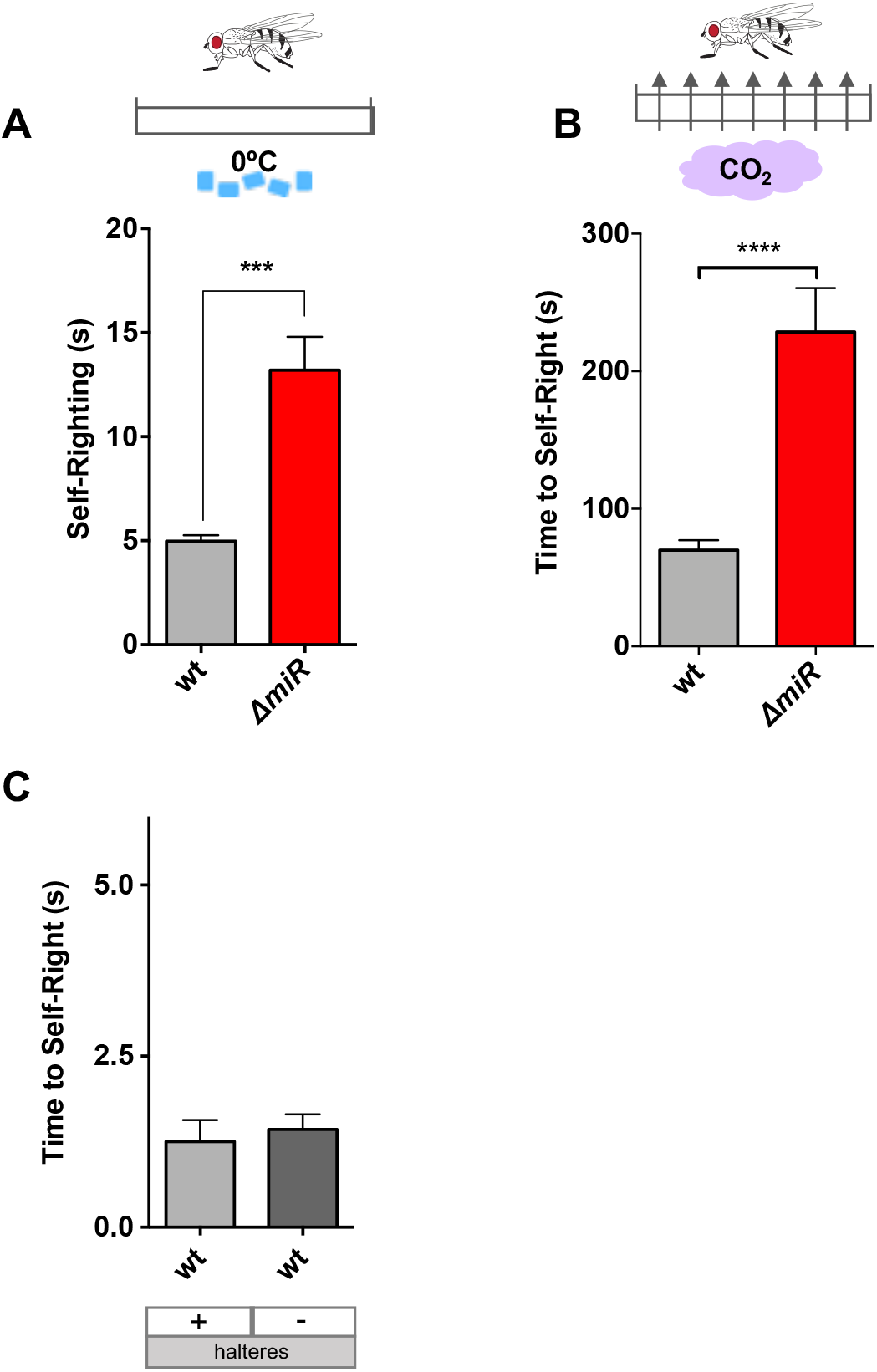
Mutation of the miRNA locus *miR-iab4/8* leads to self-righting (SR) defects in the *Drosophila* adult. Quantification of the ability of wild type (grey) and *miR-iab4/8* mutant flies (red) *(DmiR)* to return to normal orientation when turned upside down (self-righting, SR) shows significant effects of the *miR-iab4/8* locus on adult SR when tested in different experimental conditions. **A**. Ice anaesthesia. Prior to the experiment flies were maintained on ice (0°C) for 10 minutes to allow subject manipulation (mean ± SEM; n = 21-45). **B**. CO_2_ anaesthesia. Prior to the experiment flies were anaesthetised by brief exposure to CO_2_ (mean ± SEM; n = 29-44). A nonparametric Mann-Whitney U test was performed to compare treatments; p < 0.001 (***). **C**. No detectable role of halteres in adult SR. Comparison of flies with and without halteres shows that surgical removal of halteres leads to no statistically-significant effect of halteres in adult SR (mean ± SEM; n = 12-15).

**Figure S2.**
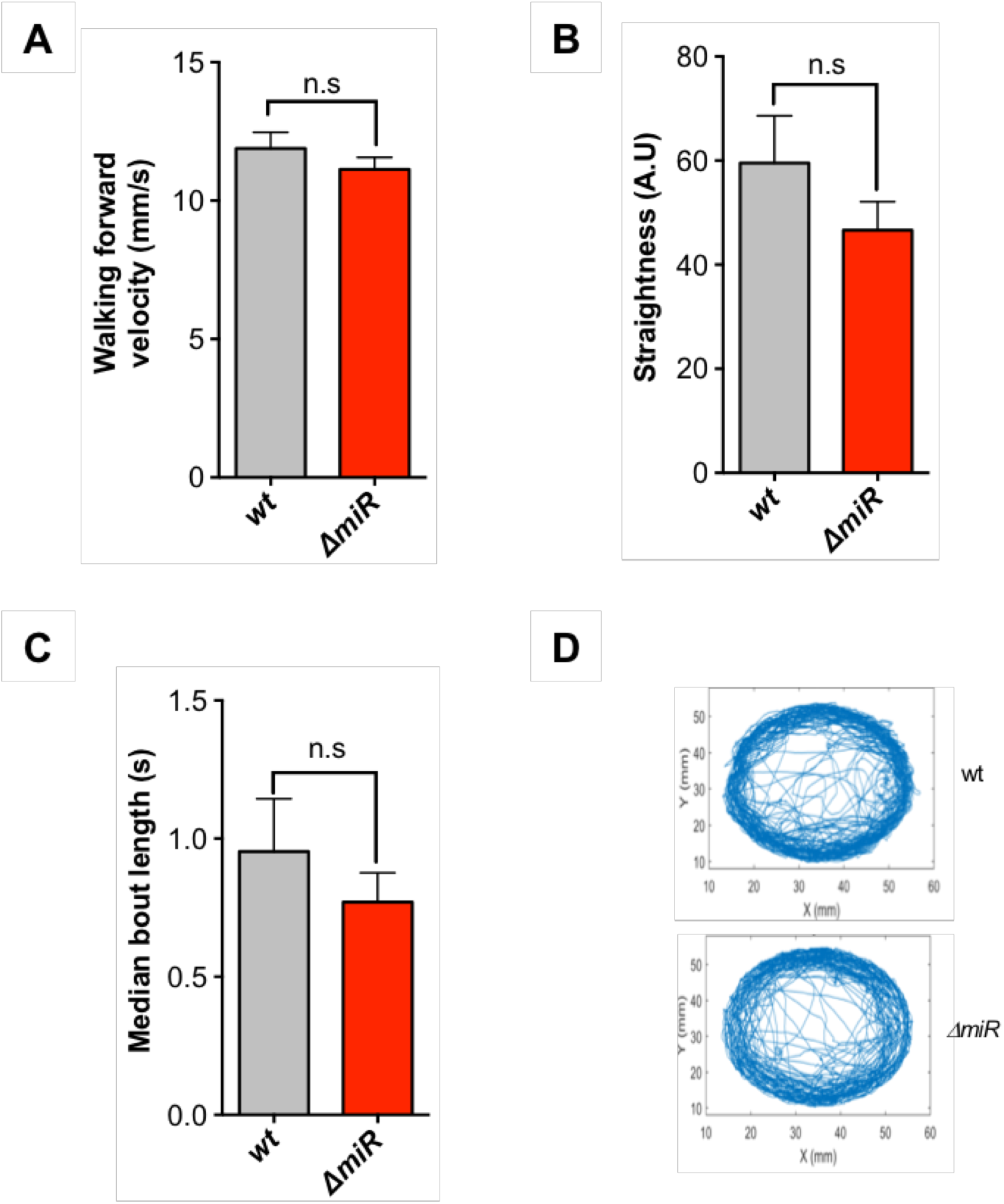
Genetic removal of *miR-iab4/8* does not affect general locomotion activities in *Drosophila* adults. Quantification of **(A)** walking velocity, **(B)** straightness, **(C)** bout length and **(D)** patterns of free walking in wild type (grey) and *miR-iab4/8* mutant flies (red) *(DmiR)* shows no statistically significant differences among the phenotypes demonstrating that absence of the *miR-iab4/8* system does not lead to a general locomotor deficit in adult flies (mean ± SEM; n = 11-20). A nonparametric Mann-Whitney U test was performed to compare treatments.

**Figure S3.**
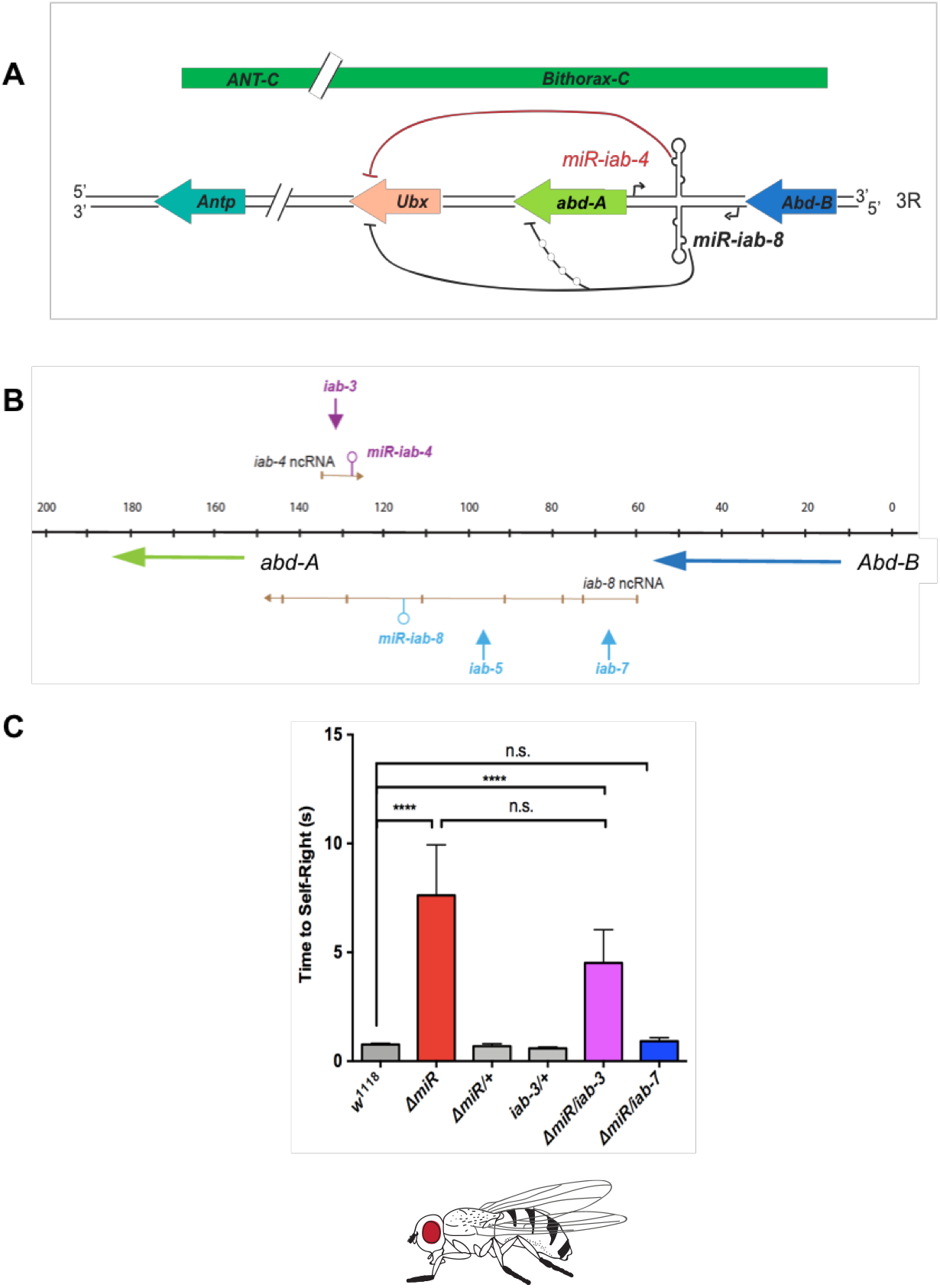
Genetic complementation tests determine that loss of *miR-iab4* leads to the adult SR phenotype. **(A)** Diagram of the *Drosophila Hox* complexes (Antennapedia (ANT-C) and Bithorax (BX-C) showing the genomic location of the *Hox* genes *Antp, Ubx, abd-A* and *Abd-B* and the miRNA system *miR-iab4/8.* Note that transcription of *miR-iab4* and *miR-iab8* occurs from opposite DNA strands. **(B)** Diagram of a sub-region of the bithorax complex showing *miR-iab-4* (magenta) and *miR-iab-8* (blue) non-coding RNAs (ncRNA), and breakpoints of rearrangement affecting *miR-iab-4* (iab-3, magenta) and *miR-iab-8* (iab-5 and iab-7, blue). **(C)**. Genetic complementation using trans-heterozygote flies for *ΔmiR* and a series of chromosomal rearrangement breakpoints *(iab-3* and *iab-5* or *iab-7)* establish that *miR-iab-4* (and not *miR-iab8)* underlies SR effects in the adult (mean ± SEM; n = 15-50). One-way ANOVA with the post hoc Tukey-Kramer (Figure E) tests were performed to compare treatments; P > 0.05 (nonsignificant; n.s.) and p < 0.001 (***).

**Figure S4.**
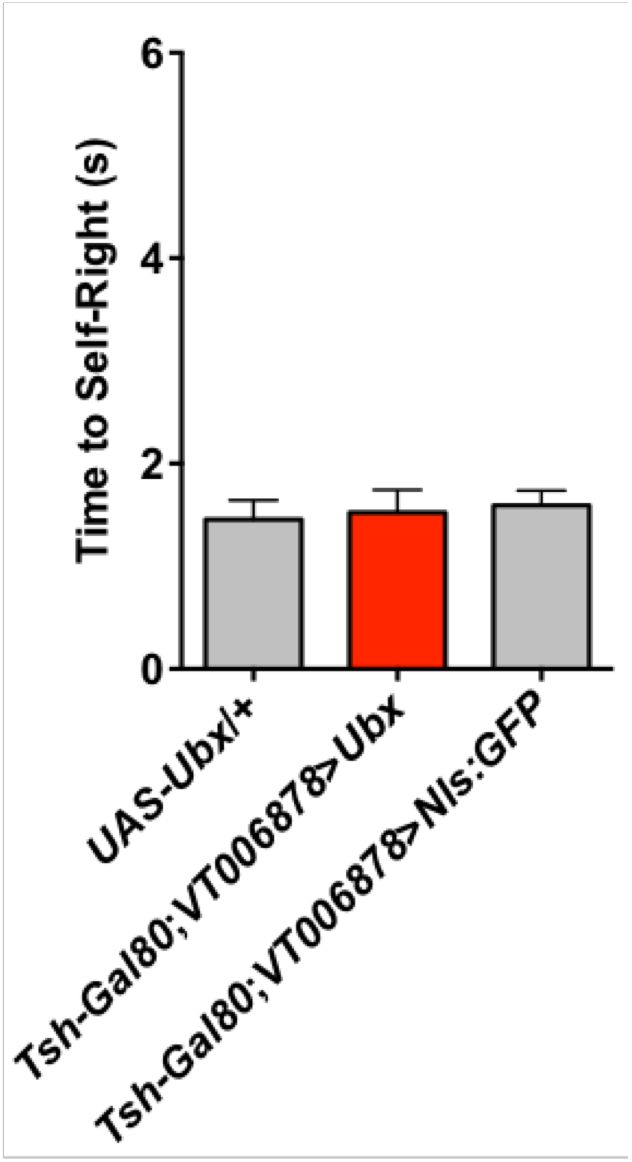
Upregulation of *Ubx* within the VT006878 domain in the brain is not sufficient to generate a self-righting (SR) phenotype. Expression of *VT006878> Ubx* (red) in the presence of *Tsh-Gal80* (which represses expression in the entire ventral nerve cord, VNC) leads to no statistically significant changes in SR times when compared with control lines *(UAS-Ubx/+)* (grey) and *Tsh-Gal80; VT006878>Ubx* (grey) (mean ± SEM; n = 22). One-way ANOVA with the post hoc Tukey-Kramer (Figure E) test were performed to compare treatments.

**Figure S5.**
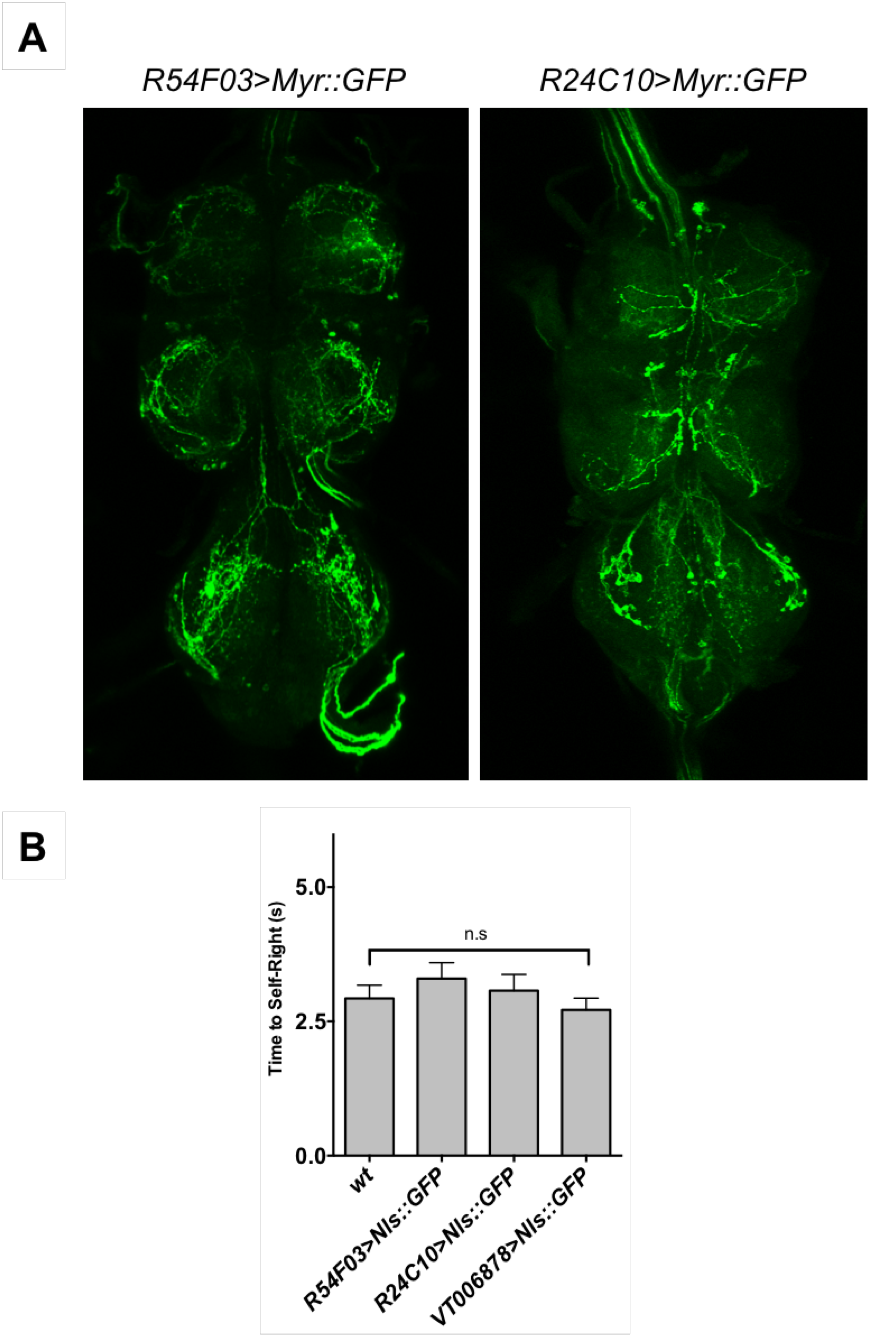
Roles of specific motor neuron subpopulations in adult SR behaviour. **A**. Patterns of motoneuronal Gal4 drivers expressing in VNC related to Figure 3. Adult VNC expression membrane-associated GFP (Myr::GFP) under control of R54F03-Gal4 *(R54F03> Myr::GFP)* or R24C10-Gal4 *(R24C10> Myr::GFP).* **B**. Flies that expressed GFP as control do not show significant changes in SR times compared with control wt (w^1118^) (mean ± SEM; n = 41). A nonparametric Mann-Whitney U test were performed to compare treatments; P > 0.05 (nonsignificant; n.s.).

**Figure S6.**
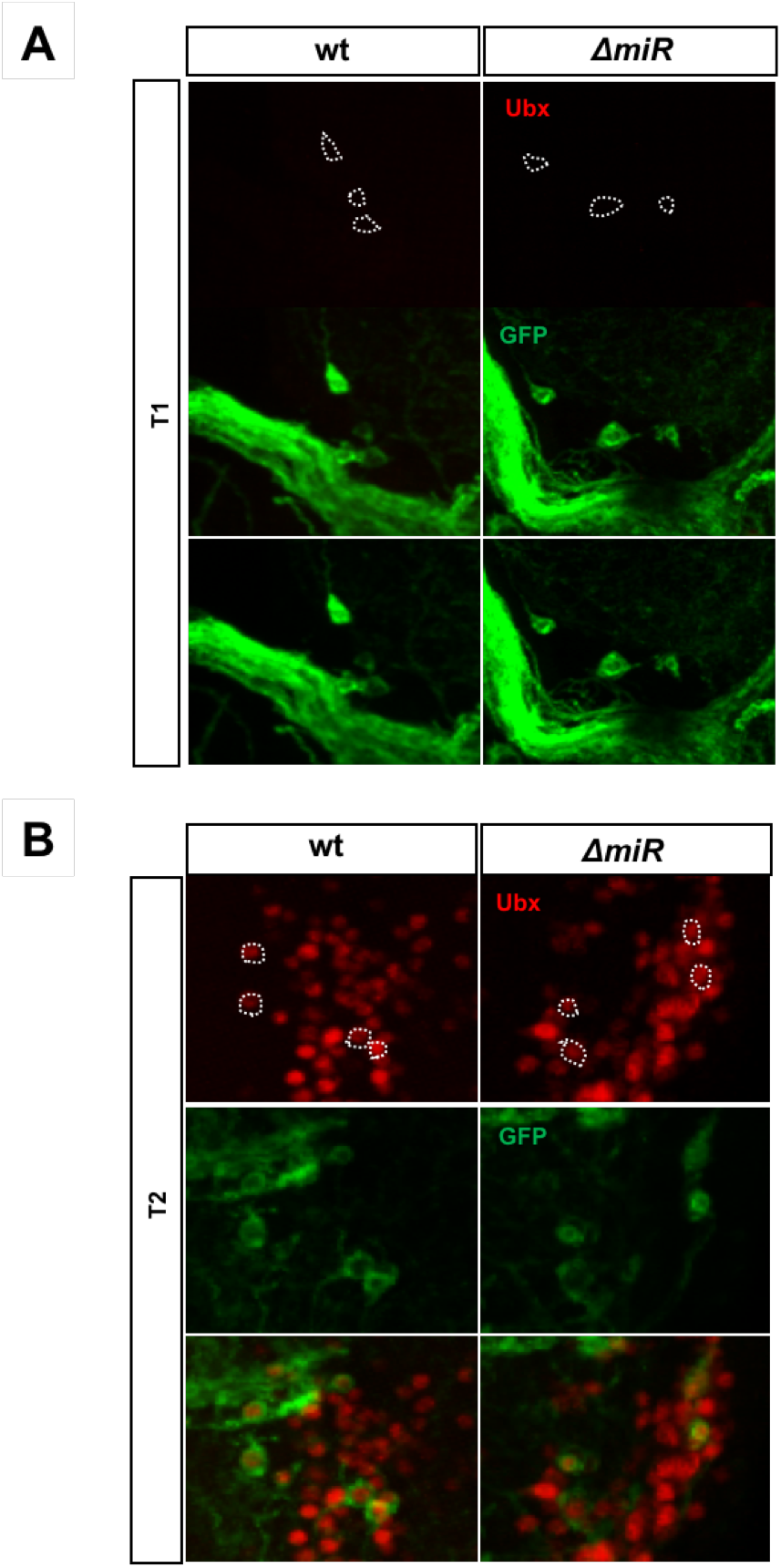
Expression pattern of Ubx protein within the VT006878 domain in the T1 and T2 ganglia of the VNC in wild type and miRNA mutants. **(A)** There is no expression of Ubx protein within the VT006878 domain in the T1 ganglion. **(B)** Expression of Ubx protein within the VT006878 domain in the T2 ganglion of normal and mutant adult flies shows no differences in expression across the genotypes (See Figure 4).

**Figure S7.**
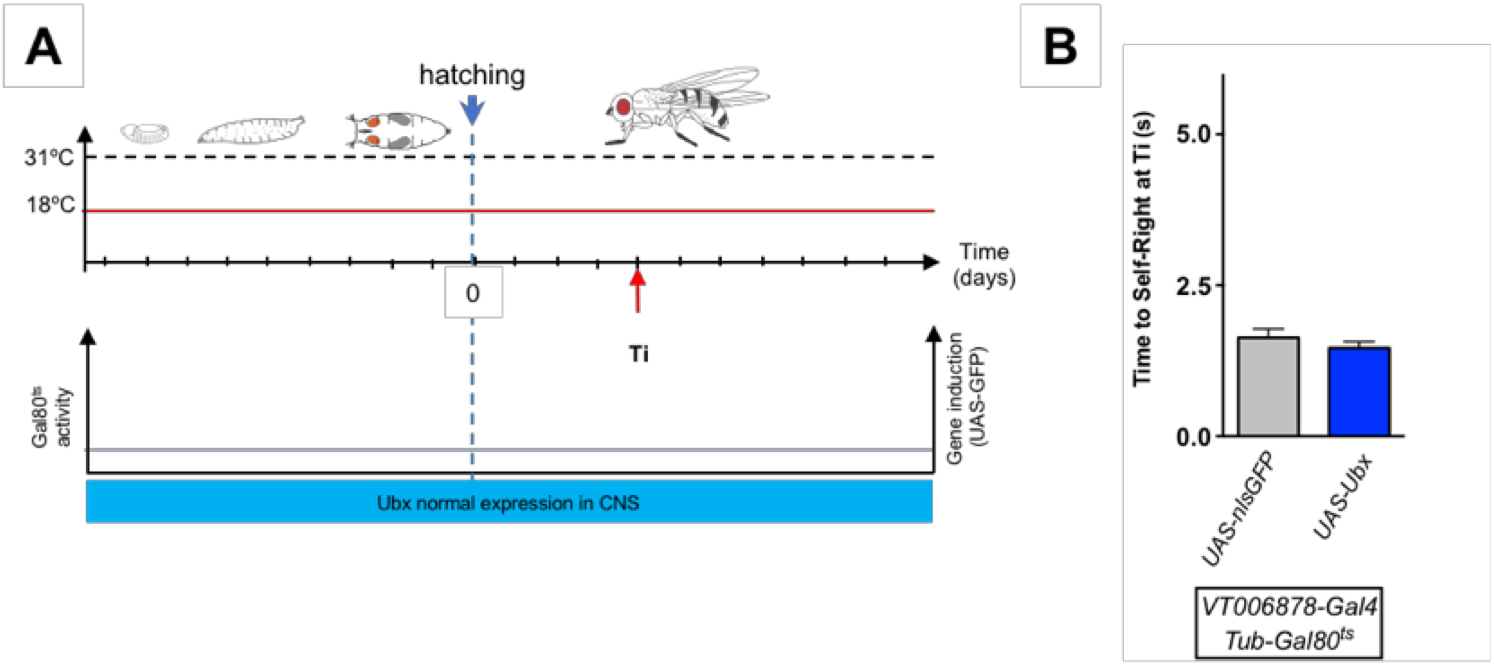
A control to the conditional expression of Ubx in adult *Drosophila*. **(A)** Graphic representation of *Gal4* and *Gal80* activities over developmental time. At 18°C, Gal80^ts^ represses Gal4 activity thus blocking *VT006878-Gal4* mediated induction of Ubx in VT006878/lin15 neurons. **(B)** Under Gal80-mediated repression, there is no induction of Ubx expression in VT006878 cells and no statistically-significant changes in SR times are observed when comparing *(VT006878;Tub-Gal80>Ubx)* (blue) with the control line *(VT006878;Tub-Gal80·>nls-GFP)* (grey) (mean ± SEM; n = 19-25). A nonparametric Mann-Whitney U test were performed to compare treatments.

## REFERENCES

Alonso, C. R. (2002). Hox proteins: sculpting body parts by activating localized cell death. Curr Biol 12, R776–8.

Ashe, V. M. (1970). The righting reflex in turtles: a description and comparison. Psychon. Sci. 20, 150–152.

Baek, M., and Mann, R.S. (2009). Lineage and birth date specify motor neuron targeting and dendritic architecture in adult *Drosophila*. J Neurosci 29, 6904 – 6916.

Bartel, D. P. (2018). Metazoan MicroRNAs. Cell 173, 20–51.

Bender, W. (2008). MicroRNAs in the Drosophila bithorax complex. Genes Dev 22, 14–19.

Benzer, S. (1967). Behavioral mutants of drosophila isolated by countercurrent distribution. Proc Natl Acad Sci U S A 58, 1112–1119.

Bodily, K. D., Morrison, C. M., Renden, R. B., and Broadie, K. (2001). A novel member of the Ig superfamily, turtle, is a CNS-specific protein required for coordinated motor control. J Neurosci 21, 3113–3125.

Branson, K., Robie, A. A., Bender, J., Perona, P., and Dickinson, M. H. (2009). High-throughput ethomics in large groups of Drosophila. Nat Methods 6, 451–457.

Bridges, C.B., and Morgan, T. H. (1923). The third-chromosome group of mutant characters of Drosophila melanogaster. Carnegie Institution of Washington 327.

Brierley, D.J., Rathore, K., VijayRaghavan, K., and Williams, D.W. (2012). Developmental origins and architecture of *Drosophila* leg motoneurons. J Comp Neurol 520,1629–49.

Carhan, A., Reeve, S., Dee, C. T., Baines, R. A., and Moffat, K. G. (2004). Mutation in slowmo causes defects in Drosophila larval locomotor behaviour. Invert Neurosci 5, 65–75.

Chen, C. L., Hermans, L., Viswanathan, M. C., Fortun, D., Aymanns, F., Unser, M., Cammarato, A., Dickinson, M. H., and Ramdya, P. (2018). Imaging neural activity in the ventral nerve cord of behaving adult Drosophila. Nat Commun 9, 4390.

Chen, T. W., Wardill, T. J., Sun, Y., Pulver, S. R., Renninger, S. L., Baohan, A., Schreiter, E. R., Kerr, R. A., Orger, M. B., Jayaraman, V., Looger, L. L., Svoboda, K., and Kim, D. S. (2013). Ultrasensitive fluorescent proteins for imaging neuronal activity. Nature 499, 295–300.

Cox, C. L., Denk, W., Tank, D. W., and Svoboda, K. (2000). Action potentials reliably invade axonal arbors of rat neocortical neurons. Proc Natl Acad Sci U S A 97, 9724–9728.

Crisp, S., Evers, J. F., Fiala, A., and Bate, M. (2008). The development of motor coordination in *Drosophila* embryos. Development 135, 3707–3717.

De Belle, J. S., Hilliker, A. J., and Sokolowski, M. B. (1989). Genetic localization of foraging (for): a major gene for larval behavior in Drosophila melanogaster. Genetics 123, 157–163.

De Navas, L., Foronda, D., Suzanne, M., and Sánchez-Herrero, E. (2006). A simple and efficient method to identify replacements of P-lacZ by P-Gal4 lines allows obtaining Gal4 insertions in the bithorax complex of Drosophila. Mech Dev 123, 860–867.

Dixit, R., Vijayraghavan, K., and Bate, M. (2008). Hox genes and the regulation of movement in Drosophila. Dev Neurobiol 68, 309–316.

Enriquez J., Venkatasubramanian L., Baek M., Peterson M., Aghayeva U., Mann R.S. (2015). Specification of individual adult motor neuron morphologies by combinatorial transcription factor codes. Neuron 86, 955–970.

Faisal, A. A. and Matheson, T. (2001). Coordinated righting behaviour in locusts. J. Exp. Biol. 204, 637–648

Hotta, Y., and Benzer, S. (1972). Mapping of behaviour in Drosophila mosaics. Nature 240,527–535.

Jusufi, A., Zeng, Y., Full, R. J. and Dudley, R. (2011). Aerial righting reflexes in flightless animals. Integr. Comp. Biol. 51, 937–943.

Karch, F., Weiffenbach, B., Peifer, M., Bender, W., Duncan, I., Celniker, S., Crosby, M., and Lewis, E. B. (1985). The abdominal region of the bithorax complex. Cell 43, 81–96.

Kitamoto, T. (2001). Conditional modification of behavior in Drosophila by targeted expression of a temperature-sensitive shibire allele in defined neurons. J Neurobiol 47, 81–92.

Lacin, H., and Truman, J. W. (2016). Lineage mapping identifies molecular and architectural similarities between the larval and adult Drosophila central nervous system. Elife 5, e13399.

Landmesser, L. T., and O’Donovan, M. J. (1984). Activation patterns of embryonic chick hind limb muscles recorded in ovo and in an isolated spinal cord preparation. J. Physiol 347, 189–204.

Lewis, E. B. (1978). A gene complex controlling segmentation in Drosophila. Nature 276, 565–570.

Mallo, M., and Alonso, C. R. (2013). The regulation of Hox gene expression during animal development. Development 140, 3951–3963.

McGinnis, W., and Krumlauf, R. (1992). Homeobox genes and axial patterning. Cell 68, 283–302.

Nalbach, G. (1993). The halteres of the blowfly. I. Kinematics and dynamics. J. Comp. Physiol. – A Sensory, Neural, Behav. Physiol. 173, 293–300.

Osborne, K. A., Robichon, A., Burgess, E., Butland, S., Shaw, R. A., Coulthard, A., Pereira, H. S., Greenspan, R. J., and Sokolowski, M. B. (1997). Natural behavior polymorphism due to a cGMP-dependent protein kinase of Drosophila. Science 277, 834–836.

Penn, D. and Brockmann, H. J. (1995). Age-biased stranding and righting in male horseshoe crabs, Limulus polyphemus. Animal Behav. 49, 1531–1539.

Petreanu, L., Gutnisky, D. A., Huber, D., Xu, N. L., O’Connor, D. H., Tian, L., Looger, L., and Svoboda, K. (2012). Activity in motor-sensory projections reveals distributed coding in somatosensation. Nature 489, 299–303.

Pfeiffer, B. D., Ngo, T. T., Hibbard, K. L., Murphy, C., Jenett, A., Truman, J. W., and Rubin, G. M. (2010). Refinement of tools for targeted gene expression in Drosophila. Genetics 186,735–755.

Picao-Osorio, J., Johnston, J., Landgraf, M., Berni, J., and Alonso, C. R. (2015). MicroRNA-encoded behavior in Drosophila. Science 350, 815–820.

Picao-Osorio, J., Lago-Baldaia, I., Patraquim, P., and Alonso, C. R. (2017). Pervasive Behavioral Effects of MicroRNA Regulation in. Genetics 206, 1535–1548.

Reed, H. C., Hoare, T., Thomsen, S., Weaver, T. A., White, R. A., Akam, M., and Alonso, C. R. (2010). Alternative splicing modulates Ubx protein function in Drosophila melanogaster. Genetics 184, 745–758.

Ronshaugen, M., Biemar, F., Piel, J., Levine, M., and Lai, E. C. (2005). The Drosophila microRNA iab-4 causes a dominant homeotic transformation of halteres to wings. Genes Dev 19, 2947–2952.

Rozowski, M., and Akam, M. (2002). Hox gene control of segment-specific bristle patterns in Drosophila. Genes Dev 16, 1150–1162.

Saint-Amant, L. and Drapeau, P. (1998). Time course of the development of motor behaviors in the zebrafish embryo. J. Neurobiol. 37, 622–632.

Sánchez-Herrero, E., Vernós, I., Marco, R., and Morata, G. (1985). Genetic organization of Drosophila bithorax complex. Nature 313, 108–113.

Seeholzer, L. F., Seppo, M., Stern, D. L., and Ruta, V. (2018). Evolution of a central neural circuit underlies Drosophila mate preferences. Nature 559, 564–569.

Shaver, S. A., Riedl, C. A., Parkes, T. L., Sokolowski, M. B., and Hilliker, A. J. (2000). Isolation of larval behavioral mutants in Drosophila melanogaster. J Neurogenet 14, 193–205.

Simon, J. C., and Dickinson, M. H. (2010). A new chamber for studying the behavior of Drosophila. PLoS One 5, e8793.

Stark, A., Bushati, N., Jan, C. H., Kheradpour, P., Hodges, E., Brennecke, J., Bartel, D. P., Cohen, S. M., and Kellis, M. (2008). A single Hox locus in Drosophila produces functional microRNAs from opposite DNA strands. Genes Dev 22, 8–13.

Suzue, T. (1996). Movements of mouse fetuses in early stages of neural development studied *in vitro*. Neurosci Lett 218, 131–134.

Truman, J. W., and Riddiford, L. M. (1999). The origins of insect metamorphosis. Nature 401,447–452.

Tyler, D. M., Okamura, K., Chung, W. J., Hagen, J. W., Berezikov, E., Hannon, G. J., and Lai, E. C. (2008). Functionally distinct regulatory RNAs generated by bidirectional transcription and processing of microRNA loci. Genes Dev 22, 26–36.

White, R. A., and Wilcox, M. (1985). Distribution of Ultrabithorax proteins in Drosophila. EMBO J 4, 2035–2043.

Yang, P., Shaver, S. A., Hilliker, A. J., and Sokolowski, M. B. (2000). Abnormal turning behavior in Drosophila larvae. Identification and molecular analysis of scribbler (sbb). Genetics 155, 1161–1174.

Zimmermann, M. J. Y., Nevala, N. E., Yoshimatsu, T., Osorio, D., Nilsson, D. E., Berens, P., and Baden, T. (2018). Zebrafish Differentially Process Color across Visual Space to Match Natural Scenes. Curr Biol 28, 2018–2032.e5.

